# *Drosophila* Sex Peptide Controls the Assembly of Lipid Microcarriers in Seminal Fluid

**DOI:** 10.1101/2020.04.24.059238

**Authors:** S. Mark Wainwright, Cláudia C. Mendes, Aashika Sekar, Benjamin Kroeger, Josephine E.E.U. Hellberg, Shih-Jung Fan, Abigail Pavey, Pauline Marie, Aaron Leiblich, Carina Gandy, Laura Corrigan, Rachel Patel, Stuart Wigby, John F. Morris, Deborah C.I. Goberdhan, Clive Wilson

**Author notes:** Corresponding Author: Clive Wilson.

## Abstract

Seminal fluid plays an essential role in promoting male reproductive success and modulating female physiology and behaviour. In the fruit fly, *Drosophila melanogaster*, Sex Peptide (SP) is the best-characterised protein mediator of these effects. It is secreted from the paired male accessory glands (AGs), which, like the mammalian prostate and seminal vesicles, generate most of the seminal fluid contents. After mating, SP binds to spermatozoa and is retained in the female sperm storage organs. It is gradually released by proteolytic cleavage and induces several long-term post-mating responses including ovulation, elevated feeding and reduced receptivity to remating, primarily signalling through the SP receptor (SPR). Here, we demonstrate a previously unsuspected SPR-independent function for SP. We show that, in the AG lumen, SP and secreted proteins with membrane-binding anchors are carried on abundant, large neutral lipid-containing microcarriers, also found in other SP-expressing *Drosophila* species. These microcarriers are transferred to females during mating, where they rapidly disassemble. Remarkably, SP is a key assembly factor for microcarriers and is also required for the female disassembly process to occur normally. Males expressing non-functional SP mutant proteins that affect SP’s binding to and release from sperm in females also do not produce normal microcarriers, suggesting that this male-specific defect contributes to the resulting widespread defects in ejaculate function. Our data therefore reveal a novel role for SP in formation of seminal macromolecular assemblies, which may explain the presence of SP in *Drosophila* species, which lack the signalling functions seen in *D. melanogaster*.

**Significance Statement:** Seminal fluid plays a critical role in reprogramming female physiology and behaviour to promote male reproductive success. We show in the fruit fly that specific seminal proteins, including the archetypal ‘female-reprogramming’ molecule Sex Peptide, are stored in male seminal secretions in association with large neutral lipid-containing microcarriers, which rapidly disperse in females. Related structures are also observed in other Sex Peptide-expressing *Drosophila* species. Males lacking Sex Peptide have structurally defective microcarriers, leading to abnormal cargo loading and transfer to females. Our data reveal that this key signalling molecule in *Drosophila* seminal fluid is also a microcarrier assembly factor that controls transfer of other seminal factors, and that this may be a more evolutionarily ancient role of this protein.

## Introduction

In addition to spermatozoa, semen contains a complex cocktail of macromolecules and nutrients secreted by the accessory glands of the male reproductive tract. In humans, seminal plasma nutrients include fructose from the seminal vesicles and triglycerides, both major energy sources for sperm in the female (1). In addition, enzymes, such as proteases and lipases, non-enzymatic binding proteins, like lectins and cysteine-rich secretory proteins (CRISPs), and a wide range of hormones and signalling molecules are major components, many of them generated in the prostate gland (2, 3). These molecules may be stored for days in the gland following cellular secretion, prior to being delivered to females during mating, when mixing of seminal plasma components can trigger enzyme and signal activation (4). However, the mechanisms that underpin these storage and activation events are generally not well understood.

The paired *Drosophila melanogaster* male accessory glands (AGs) share functional similarities with both the prostate and seminal vesicles in humans (5). The monolayer epithelium of these glands is formed from two secretory cell types, about 1000 main cells and 40 secondary cells at the distal tip (6)(Fig. 1A, A’). This glandular epithelial tube surrounds a large lumen. The AG secretome and its functions have been extensively characterised and multiple bioactive Accessory gland proteins (Acps) identified (7, 8). Several of these induce behavioural and physiological changes in mated females. The archetypal Acp is Sex Peptide (or Acp70Aa), a 36 amino acid protein, which is synthesised by main cells (9, 10). On transfer to females following mating, SP effects a comprehensive reprogramming of female physiology and behaviour. It promotes long-term increases in egg-laying, reduces female receptivity to remating (11, 12) and affects sperm release (13), diet (14), feeding behaviour (15), water balance (16), defaecation (17), sleep (18), immunity (19), aggression (20) and memory (21).

**Fig. 1.**
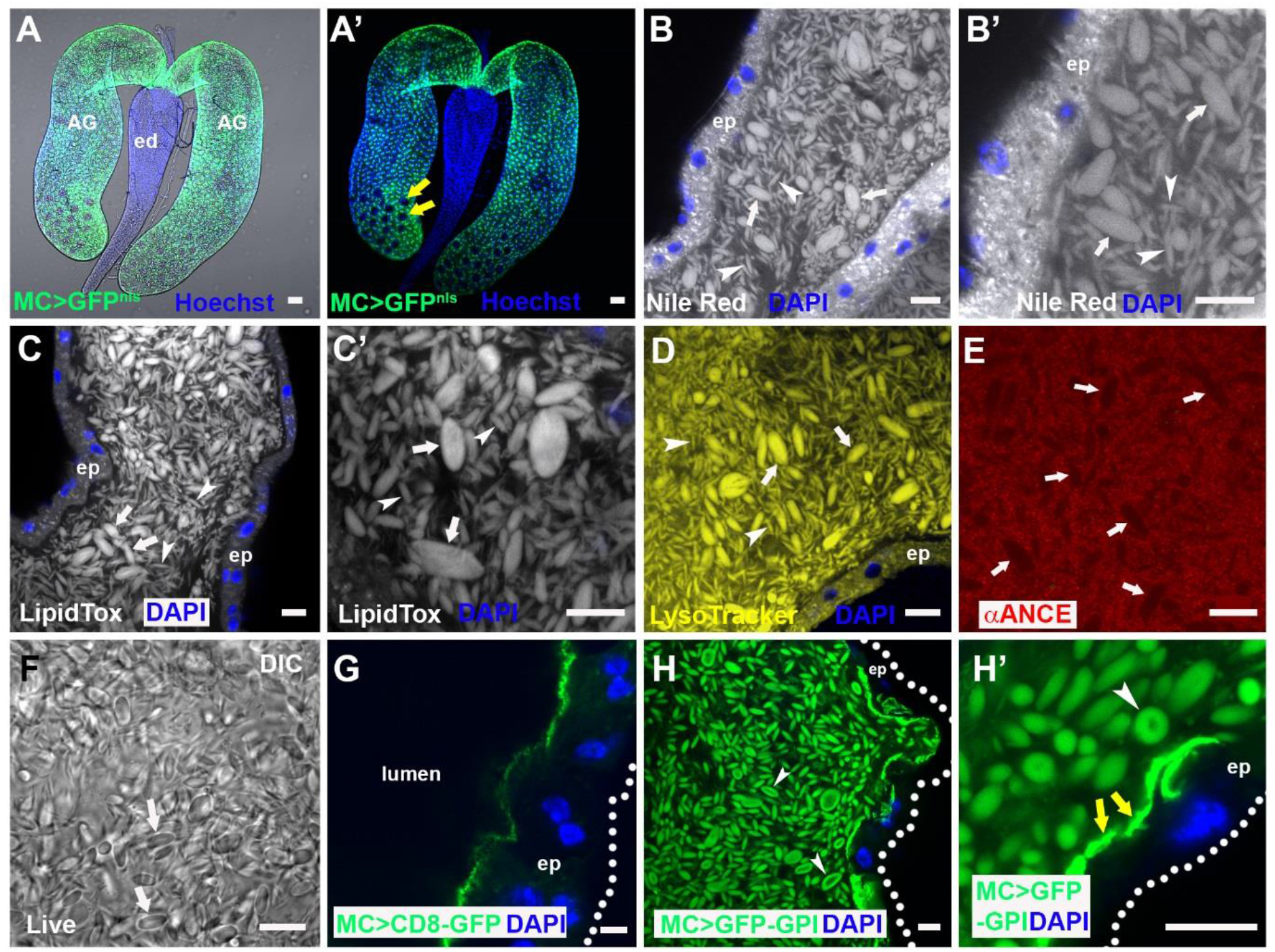
The accessory gland lumen contains abundant lipophilic microcarriers. (A, A’) Fluorescence image with (A) and without (A’) bright-field illumination of paired *Drosophila melanogaster* male accessory glands (AGs) connecting to the ejaculatory duct (ed). Main cells express nuclear GFP under *Acp26Aa*-GAL4 main cell-specific control (green), but secondary cells in distal tip (two of which are marked by yellow arrows in A’) do not. (B-E) Confocal sections through AG lumen stained with Nile Red (B, B’; latter is high magnification view), LipidTox (C, C’), LysoTracker Deep Red (D; yellow) and anti-ANCE (red), a soluble secreted protein (E). White arrows mark representative large microcarriers and arrowheads mark small microcarriers. (F) DIC image of lumen from living AG also reveals microcarriers (white arrows). (G) Transmembrane CD8-GFP expressed in main cells marks the apical plasma membrane, but not luminal microcarriers. (H, H’) Main cell-expressed GFP-GPI labels microcarriers at their surface (H, H’; white arrowheads) together with the apical surface of the epithelial monolayer (H’; yellow arrows). Nuclei marked with Hoechst (A, A’; blue) or DAPI (B-E, G, H; blue). AG epithelium (ep) (dotted white line in G, H marks approximate basal surface). Scale bars, 10 μm.

Maintaining this complex post-mating response (PMR) requires SP association with the sperm plasma membrane after mating (12). Sperm can then be stored for several weeks in two female organs, the paired spermathecae and the seminal receptacle, with SP gradually released by proteolytic cleavage to mediate its effects (22).

Studies in which SP or SP mutant peptides are either injected or expressed ectopically in females have demonstrated that SP can induce many of the characterised female PMRs, with distinct molecular domains in SP having different functions eg., (9, 23, 24). In females, the SP receptor (SPR) is required to mediate most of these effects (25). SPR is expressed in specific neurons of the female reproductive tract that have a key role in the SP-dependent PMR (26, 27), and in other neurons in the CNS that are also able to respond to circulating SP (28, 21). In addition, SP appears to produce some SPR-independent post-mating responses in females (29, 20).

Here we report a novel SPR-independent function for SP in males, involving storage and delivery of seminal components. We show that the AG lumen is filled with many large, fusiform- and ellipsoid-shaped microcarriers containing a neutral lipid core and coated with specific proteins such as SP. Microcarriers rapidly dissipate on transfer to females after mating, providing a simple mechanism for timely release of stored seminal proteins. Surprisingly, we find that SP is essential for assembly of microcarriers in males, and that this function is required for the normal delivery of microcarrier-associated macromolecules and nutrients to the female reproductive tract during mating. Furthermore, we identify related microcarrier structures in other *Drosophila* species that express a Sex Peptide and show that the size and shape of microcarriers has changed as the amino acid sequence of Sex Peptide evolved in these species.

## Results

### The lumen of the accessory gland is filled with large neutral lipid-containing microcarriers

While analysing the lipid content of epithelial cells within the male AG, using the lipophilic dye Nile Red, which stains membranes and lipid droplets, we observed that the large AG lumen is filled with fluorescent fusiform structures typically 3-8 μm in length (Fig. 1B, B’). These structures were of variable diameter ranging from less than 0.5 μm to a maximum of 4.0 μm (Fig. S1I). Since these structures were found to bind specific main cell proteins (Fig. 2A), we call them ‘microcarriers’. The neutral lipid-specific dye LipidTox Red stained microcarriers highly selectively (Fig. 1C, C’), suggesting they contain large quantities of triglycerides and other non-polar lipids. Microcarriers were also detected using high concentrations of the acidophilic, but partially hydrophobic, LysoTracker dyes (Fig. 1D; (30)). By contrast, in fixed tissue, microcarriers exclude access to antibodies raised against soluble secreted AG proteins, such as angiotensin I-converting enzyme (ANCE; Fig. 1E). Microcarriers are not an artefact of fixation or staining, because they are readily discernable in living glands using Differential Interference Contrast (DIC) microscopy (Fig. 1F).

**Fig. 2.**
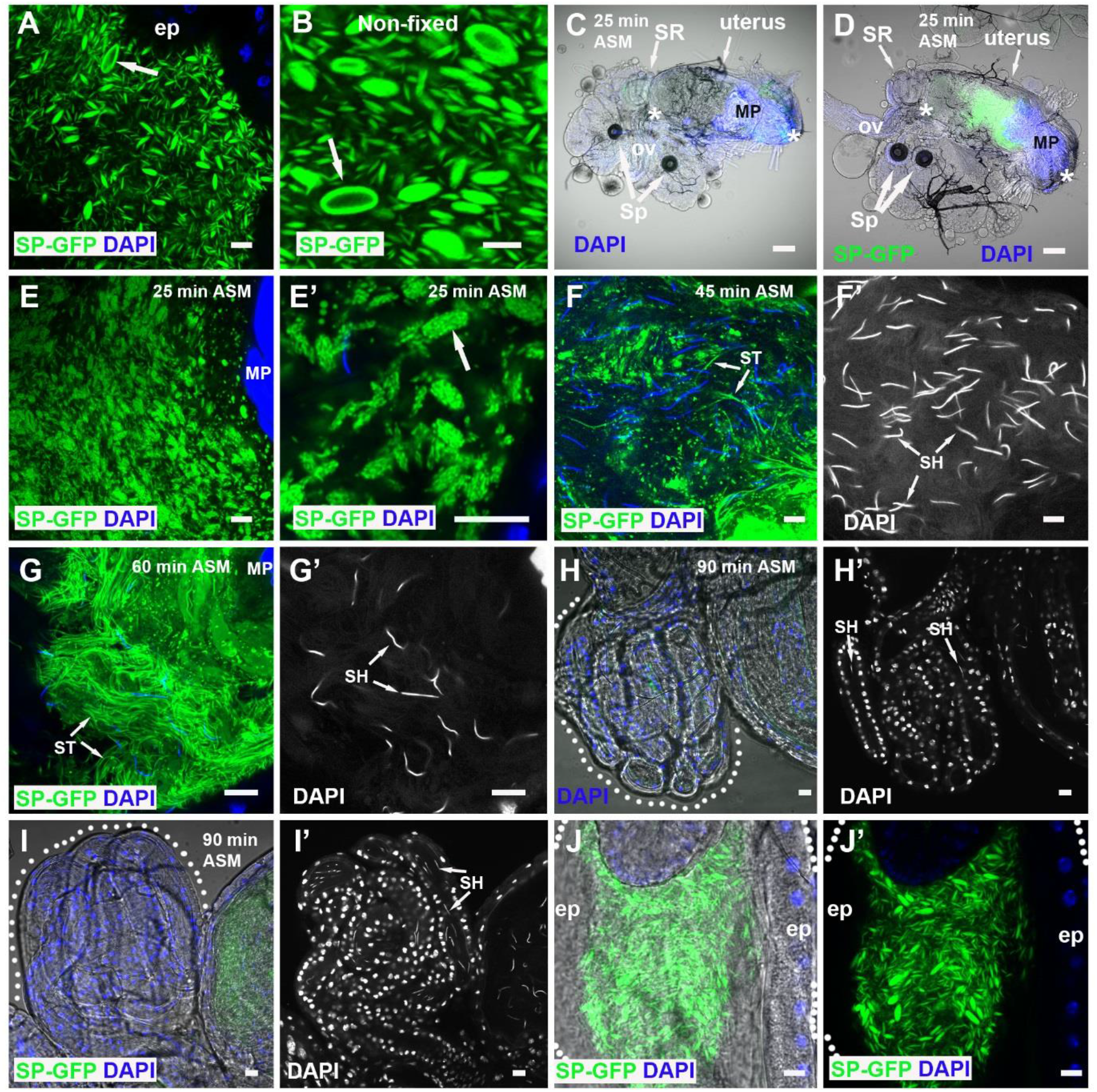
SP-GFP is loaded on microcarriers, which disassemble when transferred to females. (A, B) SP-GFP (green) marks microcarriers in fixed (A) and non-fixed (B) AG lumen, coating the surface of the largest structures (arrows). (C, D) Combined fluorescence and bright-field images of reproductive tract of female mated to a control (C) or SP-GFP (D) male dissected 25 min after start of mating (ASM). Anterior (left) and posterior (right) limits of uterus are demarcated by white asterisks, and seminal receptacle (SR), paired spermathecae (Sp), common oviduct (ov) and mating plug (MP), which autofluoresces in the DAPI channel, are marked. (E, F) Higher magnification views of posterior uterus at this time reveal microcarrier structures have changed (E) with SP-GFP concentrated in microdomains (E’; arrow). (F, G) Later (45 min ASM), many microcarriers have disassembled and some SP-GFP has associated with sperm tails (ST; F, F’), while later still (60 min ASM), few recognisable microcarriers remain and many more strongly labelled sperm tails are observed in the anterior uterus (ST G, G’). Sperm heads (SH) are marked by DAPI. (H, I) Labelled sperm tails (ST) are not present in the seminal receptacles (60-90 min ASM), which contain sperm heads (SH), both in females mated with control (H, H’) and SP-GFP males (I, I’). (J, J’) Microcarriers remaining in the ejaculatory duct after mating maintain their structure. Outlines of seminal receptacles (H, I) and ejaculatory duct (J, J’) are marked by dotted lines. Nuclei marked with DAPI (blue). AG and ejaculatory duct epithelia (ep). Scale bars, 10 μm except C, D, 30 μm.

Previous studies have shown that some transmembrane proteins expressed in epithelial secondary cells of the AG are secreted on exosomes (31). When transmembrane proteins were expressed in main cells, they did not associate with microcarriers (Fig. 1G, S1A, A’) and neither did dyes like PKH26 that bind to lipid bilayers (Fig. S1B, B’). A secreted form of GFP, comprised of the SP signal sequence fused to GFP (24), also failed to preferentially bind to microcarriers (Fig. S1C). By contrast, GFP-GPI, a GFP fusion protein carrying the lipid anchor glycosylphosphatidylinositol, strongly labelled microcarriers when expressed in main cells (Fig. 1H, H’), but not when made in secondary cells (Fig. S1D, D’; (32)), consistent with the idea that main cells produce these structures. Indeed, a concentrated layer of GFP-GPI-positive staining was observed at the apical surface of main cells overexpressing this transgene (Fig. 1H’), reflecting the shedding of lipophilic material from these cells. In the largest microcarriers, GFP-GPI, unlike LipidTox staining (Fig. 1C, C’), was surface-localised (Fig. 1H, H’), suggesting that these structures have a distinct outer layer, most likely a phospholipid monolayer into which the GPI anchor is inserted, surrounding the neutral lipid core. Although microcarrier ultrastructure was difficult to preserve for transmission electron microscopy, micrographs were consistent with these structures having a homogeneous internal structure (Fig. S1E).

### Sex Peptide is a microcarrier cargo

An SP-GFP C-terminal GFP fusion protein expressed in main cells under the control of *SP* gene regulatory elements (33) has previously been used to assess SP transfer to females. Surprisingly, we found that it strongly associates with microcarriers and concentrates at the surface of the largest structures, but is present at very low levels within main cells (Fig. 2A, B; SI Movie 1). When SP-GFP males were mated, fluorescently labelled microcarriers were transferred to females (Fig. 2C, D). We were unable to detect microcarriers using neutral lipid stains in the female reproductive tract, at least partly because of poor dye penetration. However, using SP-GFP as a marker, we found that 25 min after the start of mating (ASM), which is typically within five minutes of the end of mating, microcarriers had already started to change their morphology (Fig. 2E, E’). Although their basic fusiform shape was frequently still distinguishable, SP-GFP concentrated in microdomains on the microcarrier surface. Later, at 45 min ASM, smaller spherical SP-GFP-positive puncta were dispersed throughout the uterus and SP-GFP was observed on a subset of sperm tails (Fig. 2F, F’), while later still (60 min ASM), more of the SP-GFP (Fig. 2G, G’) was associated with sperm tails. However, at 90 min ASM, only very weak, if any, GFP expression was observed on sperm in the sperm storage organs (Fig. 2H-I’), either because the most strongly labelled sperm do not migrate to these organs or because the GFP tag is lost over time. Microcarriers that are ejected from the AG, but remain in the male ejaculatory duct after mating, do not break down (Fig. 2J, J’), suggesting that microcarrier dissipation is triggered by physical or chemical signals inside the uterus.

To confirm that C-terminal tagging of SP with GFP does not affect SP’s binding properties in the AG lumen and to begin to dissect out what domains in SP bind to microcarriers, we overexpressed three SP-GFP fusions in main cells under GAL4/UAS control: the N-terminal half of mature SP fused at its C-terminus to GFP (SPn-GFP), the C-terminal half of SP fused at its N-terminus to GFP (GFP-SPc), and a fusion with GFP located in the centre of the SP protein (SPn-GFP-SPc). The latter has been shown to have biological activity in females (24). Using the main cell-specific *Acp26Aa*-GAL4 driver (Fig. 1A) (11), which expresses at lower levels than GFP-tagged SP under its own promoter, all SP fusions partitioned with microcarriers (Fig. S1F-H), albeit less selectively for the N-terminal SP construct, SPn-GFP. Microcarriers therefore appear to bind to both the N- and C-terminal domains of SP. We conclude that they act as stores for SP, other seminal proteins, such as those with a GPI anchor, and neutral lipids in males, and serve as vehicles for their transfer to females. Regulated microcarrier disassembly in females presumably assists in the timely release of lipids and seminal proteins, such as SP, after mating.

### SP controls microcarrier morphology via an SPR-independent mechanism

To assess whether *SP* is involved in microcarrier assembly, we analysed AGs of males carrying the previously generated *SP^0^* null allele (12), either as a homozygote or in transheterozygous combination with a small *SP* deficiency, *Df(3L)Δ130* (35, 12). These transheterozygous *SP* null males have been used to characterise the full range of *SP* mutant PMR phenotypes (12–21). Unexpectedly, these mutant animals displayed dramatic defects in microcarrier morphology (Fig. 3A-D, I-J; S2A, B; S3A). Most microcarrier-like structures were highly enlarged, and either spherical or ellipsoid in shape. The enlarged microcarrier phenotype was never observed in wild type glands (Fig. 3A, C). In confocal images of the AG lumen, 10/10 *SP^0^/Df(SP)* null glands had microcarriers with a minimum width greater than 10 μm, whereas 0/10 wild type glands contained such structures (*P*<0.0001; Fisher’s exact test). The enlarged microcarriers from the *SP^0^/Df(SP)* null glands were uniformly stained by LipidTox. They appeared like large lipid droplets under DIC (Fig. 3D). The defects were absent in *SP^0^ SP^+^/Df(3L)Δ130* males, which express an *SP* genomic rescue construct that rescues the PMR phenotypes in mated females (12) (Fig. 3E); 0/10 *SP* rescue glands had microcarriers with a minimum width greater than 10 μm (*P*<0.0001 versus *SP^0^/Df(SP)*). Automated measurement of minimum microcarrier diameter in individual images of male AGs with these different genotypes confirmed the change in size distribution in the *SP* null background (Fig. 3I). Mating *SP^0^/Df(SP)* null males multiple times with females over several days to mix and eject the AG’s contents, exacerbated the mutant phenotype, with some microcarriers spanning the entire diameter of the AG lumen (Fig. 3F, G), suggesting that microcarriers can enlarge by fusion. When seminal fluid remained in the ejaculatory duct of *SP^0^/Df(SP)* null males after mating, the duct lumen was also filled with enlarged microcarriers (Fig. 3H).

**Fig. 3.**
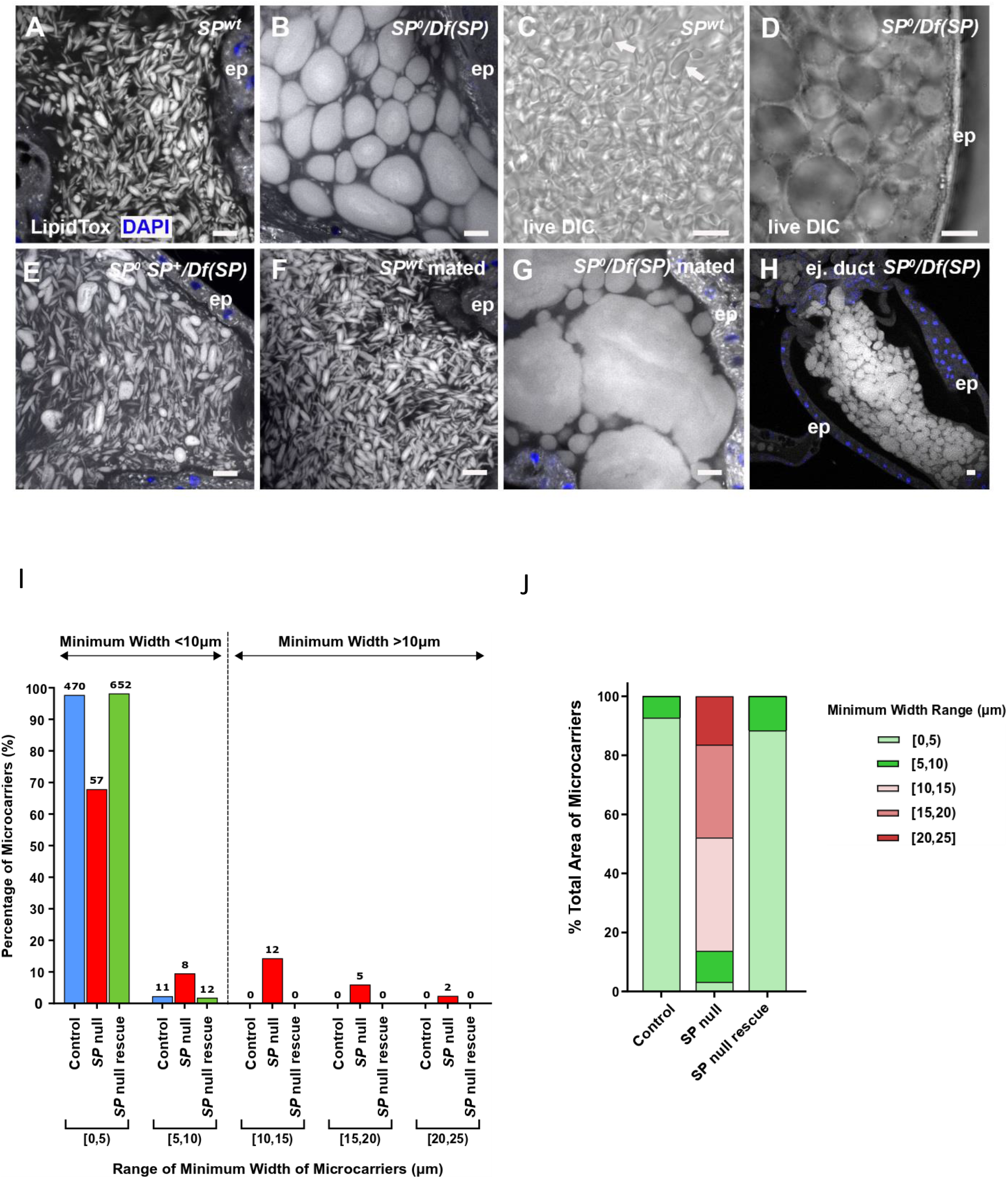
SP is essential for proper assembly of microcarriers. (A, B) Confocal images of LipidTox-labelled microcarriers in lumen of AG from control (A) and *SP^0^/Df(SP)* null (B) males. Mutant male has grossly enlarged microcarriers. (C, D) DIC images of living AGs dissected from control (C, white arrows) and *SP^0^/Df(SP)* (D) males. (E) Microcarrier structural defects in *SP^0^/Df(SP)* null males are rescued by a genomic *SP* construct *SP^0^ SP^+^/Df(SP)*. (F, G) Microcarriers enlarge further after multiple matings in *SP^0^/Df(SP)* null (G), but not in wild type (F) males. (H) In *SP^0^/Df(SP)* null males, these enlarged microcarriers are observed when seminal fluid remains in the lumen of the ejaculatory duct after mating. (I, J) Microcarrier size and area profiles for glands shown in A, B, E. Microcarrier outlines were detected in images of the AG lumen using CellProfiler Software version 2.2.0 (see Methods) and then grouped according to minimum width range (I) or percentage of luminal area occupied by microcarriers in each width range (J). Numbers of microcarriers within each size range are shown above bars (I). *SP^0^/Df(SP)* null glands have considerably fewer small microcarriers (<10 μm) and more large microcarriers (>10 μm) than the other genotypes. The enlarged microcarriers in *SP^0^/Df(SP)* null glands contain most of the lipid in the AG lumen, as estimated by LipidTox staining area. Nuclei marked with DAPI (blue). AG (A-G) or ejaculatory duct (H) epithelium (ep). Scale bars, 10 μm.

To confirm that SP expression in main cells is required for normal microcarrier assembly, we knocked down *SP* transcripts specifically in these cells, using the GAL4/UAS system (35), employing the *Acp26Aa*-GAL4 driver (11). Although limited effects were observed when these experiments were performed at 25°C, expression of *SP-*RNAis from three different transgenic lines at 29°C, a temperature that typically enhances GAL4-induced expression (36), produced consistent marked defects in microcarrier morphology (Fig. 4A-C, Fig. S2C, D, Fig. S3B). Microcarriers were enlarged in all three knockdowns, though to a lesser extent than in *SP^0^/Df(SP)* null males. As observed in mated *SP^0^/Df(SP)* null males (Fig. 3G), mating greatly exacerbated the size phenotype (Fig. 4E, F).

**Fig. 4.**
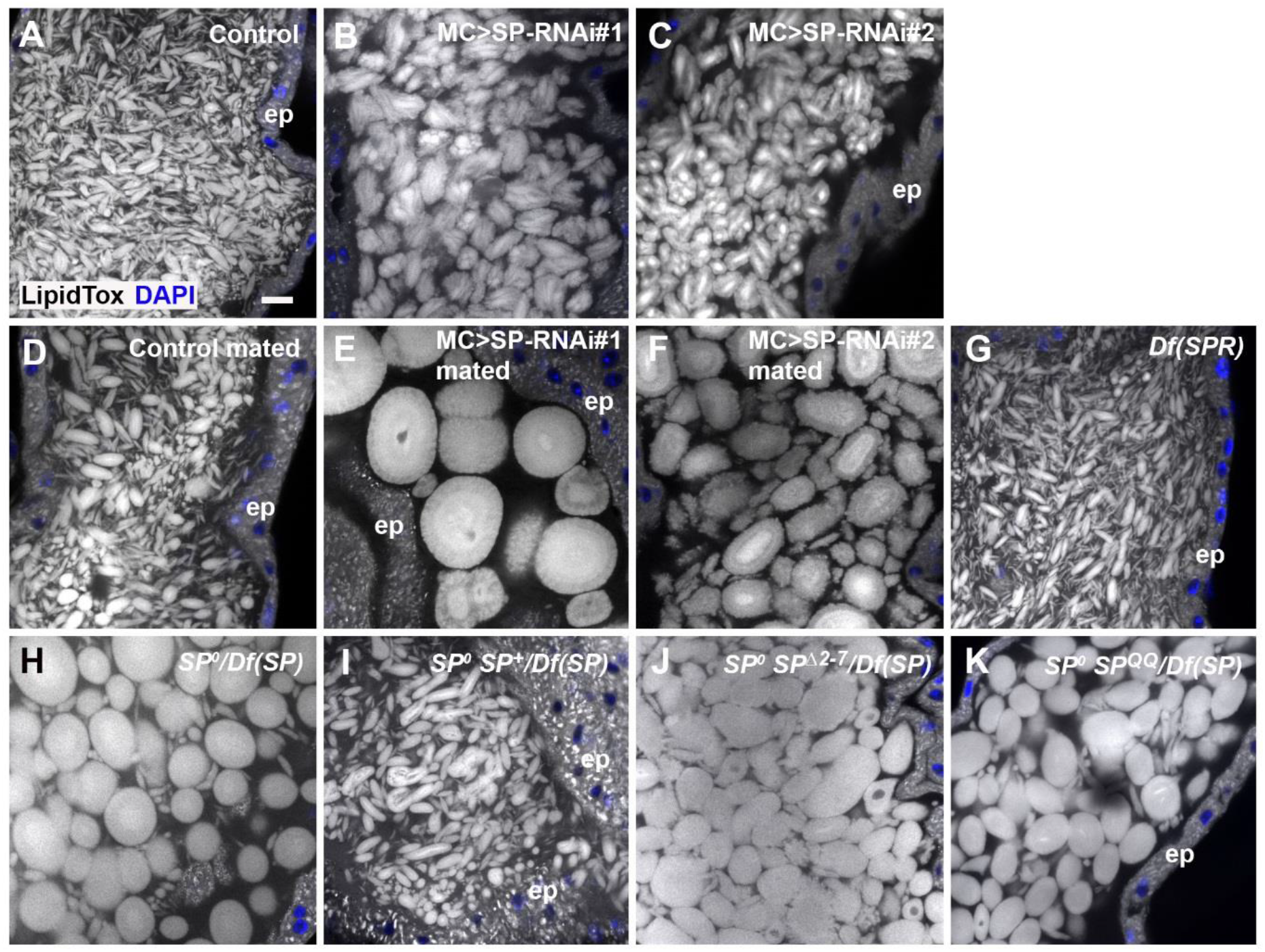
Knockdown of *SP* in main cells also produces highly enlarged microcarriers. All specimens are stained with LipidTox. (A) Confocal image of microcarriers in lumen of AG from control male. (B-C) Knockdown of *SP* in main cells at 29°C with two RNAis, *UAS-SP-RNAi#1* (B; IR2 from (11)) and *UAS-SP-RNAi#2* (C; TRiP.JF02022) produces enlarged microcarriers. (D-F) Multiple mating of *SP* knockdown males leads to further increases in microcarrier size (E, F), presumably via fusion, which is not observed in controls (D). (G) *SPR* mutant males (homozygous *Df(1)Exel6234*) have normal microcarriers. (H-K) The *SP^0^/Df(SP)* null phenotype (H) is rescued by a wild type *SP* genomic construct in *SP^0^ SP^+^/Df(SP)* males (I), but not by genomic constructs expressing mutant SP^Δ2-7^ (J) or SP^QQ^ (K). Nuclei marked with DAPI (blue). AG epithelium (ep). Scale bar in (A), 10 μm applies to all panels.

In females, many of SP’s activities in modulating the female PMR are mediated by the SPR (25). However, *SPR* mutant males displayed completely normal microcarriers (Fig. 4G), demonstrating that SP acts independently of the SPR in the male AG, presumably via direct interaction with microcarriers.

Binding of SP to the plasma membrane of sperm in females requires a short peptide sequence at the N-terminal end of the mature molecule (22). This region must be proteolytically removed for SP to be released from sperm in the sperm storage organs. Two mutants expressed under the control of the SP promoter, one that lacks the N-terminal membrane-association domain (SP^Δ2-7^) and the other mutated at the proteolytic cleavage site (SP^QQ^), have both previously been shown to fail to induce the long-term PMR in females (22). These constructs also failed to rescue the microcarrier phenotype in *SP^0^/Df(SP)* null males (Fig. 4H-K; Fig S3C). For both mutants, 7/7 glands had microcarriers with a minimum width greater than 10 μm (Fig. S3C), suggesting that the N-terminal region of SP, which appears to bind microcarriers (Fig. S1F), plays an important role in microcarrier assembly, as well as sperm binding.

### Microcarriers from *SP* mutant males do not disassemble normally in females after mating

A key property of microcarriers is that they are stable in the male AG, yet change their morphology within minutes, when transferred to females. We tested how this process is affected in *SP* mutants. Since the C-terminally tagged SP-GFP construct, which has previously been reported to lack normal SP activity in females (23), failed to rescue the *SP^0^/Df(SP)* null microcarrier phenotype in males (Fig. 5B), we used this as an alternative to neutral lipid dyes to mark microcarriers. SP-GFP distributed evenly throughout the enlarged microcarriers in *SP^0^/Df(SP)* null males (Fig. 5A, B).

**Fig. 5.**
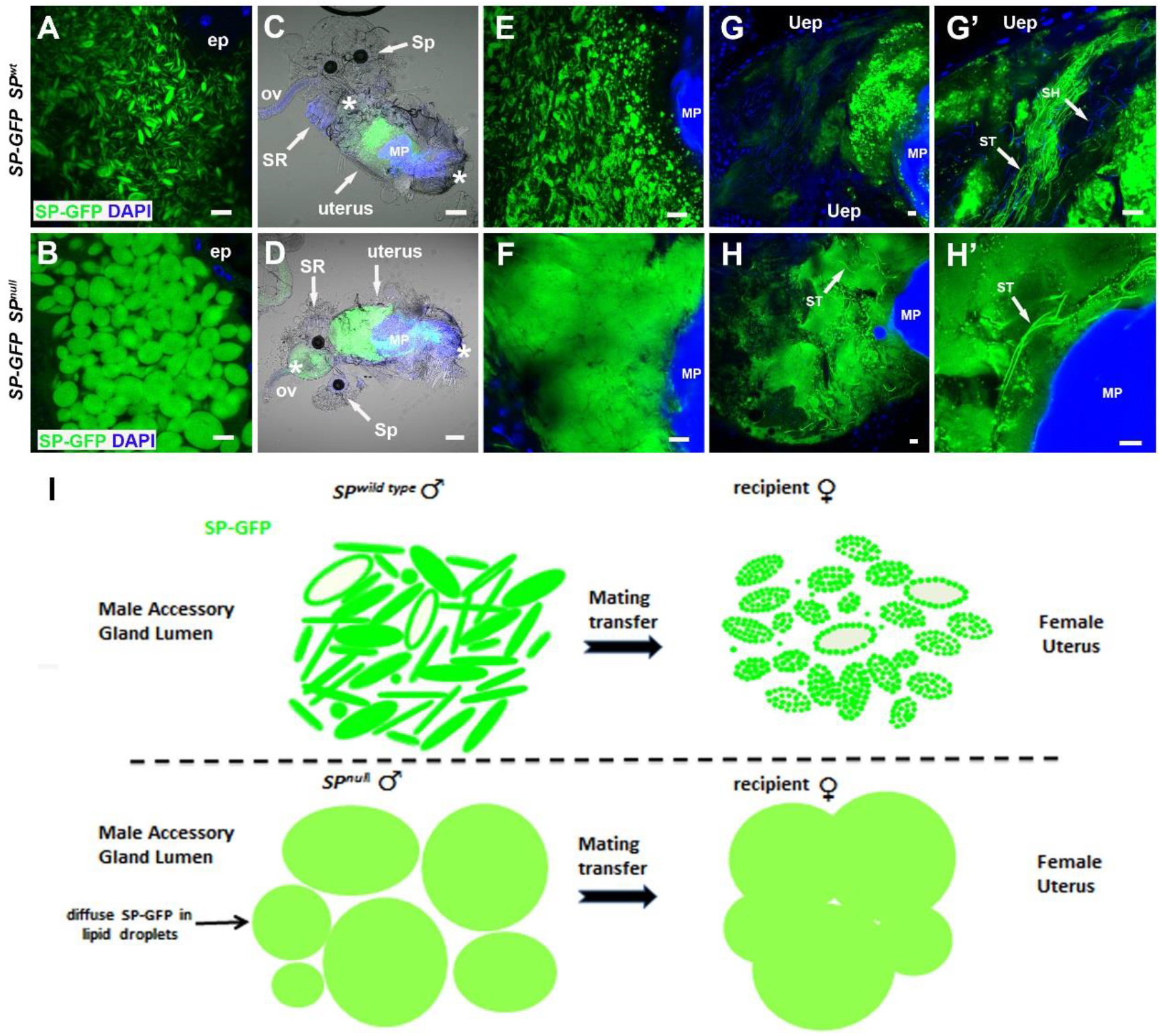
Microcarriers from *SP* null males do not dissipate normally when transferred to females during mating. (A, B) A genomic SP-GFP fusion construct labels *SP* wild-type microcarriers (A), and enlarged defective microcarriers in the AG of *SP^0^/Df(SP)* null males, though it does not rescue the associated microcarrier phenotype (B). (C-F) Combined fluorescence and bright-field images at 25-30 min ASM of whole reproductive tracts (anterior on left; C, D) and posterior uterus at higher magnification (E, F) from females mated either with control males expressing SP-GFP (C, E) or with *SP^0^/Df(SP)* null males expressing SP-GFP (D, F). Microcarrier-like structures from the *SP^0^/Df(SP)* null male are fused together in a globular mass, whereas microcarriers from control males do not fuse, but carry localised SP-GFP puncta. (G, H) At 45-50 min ASM, SP-GFP-positive material remains in a globular mass in females mated with *SP^0^/Df(SP)* null males, which extends into the anterior uterus, unlike controls (H, H’). This mass contains a few intensely labelled sperm tails (arrows). By contrast, SP-GFP from wild type males has dispersed, although some intense fluorescent puncta remain (G, G’), and often many sperm tails in the anterior uterus are labelled (arrows in G’). (I) Schematic representing microcarrier structure in accessory glands of wild-type and *SP^0^/Df(SP)* null males, as visualised using the SP-GFP fusion protein, and the changes that take place 25-30 min ASM in the female reproductive tract. In (C, D), anterior (left) and posterior (right) boundaries of uterus are demarcated by asterisks and seminal receptacle (SR), one of the two spermathecae (Sp), oviduct (ov) and mating plug (MP) are labelled. Nuclei marked with DAPI (blue). AG epithelium (ep), uterine epithelium (Uep). Scale bars, 10 μm except for C, D, 30 μm.

Unlike in controls (Fig. 5C), microcarriers from *SP^0^/Df(SP)* null males failed to rapidly dissipate in females and instead formed a homogeneously stained mass in the uterus (Fig. 5D), which did not break down during the period when SP-GFP is normally transferred to sperm tails (compare Fig. 5G, G’ with Fig. 5H, H’); indeed, unlike controls, the mass extended into the anterior uterus with some sperm tails embedded within it. We conclude that normal dissipation and distribution of microcarrier cargos is disrupted in females mated with *SP^0^/Df(SP)* null males, and this is likely to contribute to the aberrant post-mating phenotypes observed in mated females.

### SP and microcarrier structure have rapidly co-evolved in *Drosophila* species

To test whether other *Drosophilidae* might employ a similar neutral lipid-based strategy to package molecules in seminal fluid, we stained the AGs of multiple *Drosophila* species with LipidTox (Fig. 6). Species closely related to *D. melanogaster*, namely *D. simulans* and *D. sechellia* (Fig. 6A), had microcarriers with remarkably similar size and shape (Fig. 6B-D). AGs of the species, *D. yakuba*, and *D. erecta*, which are still members of the *melanogaster* group but have more divergent SP structure (24) (Fig. S4), also contained microcarriers, but these were more spherical in shape (Fig. 6E, F).

**Fig. 6.**
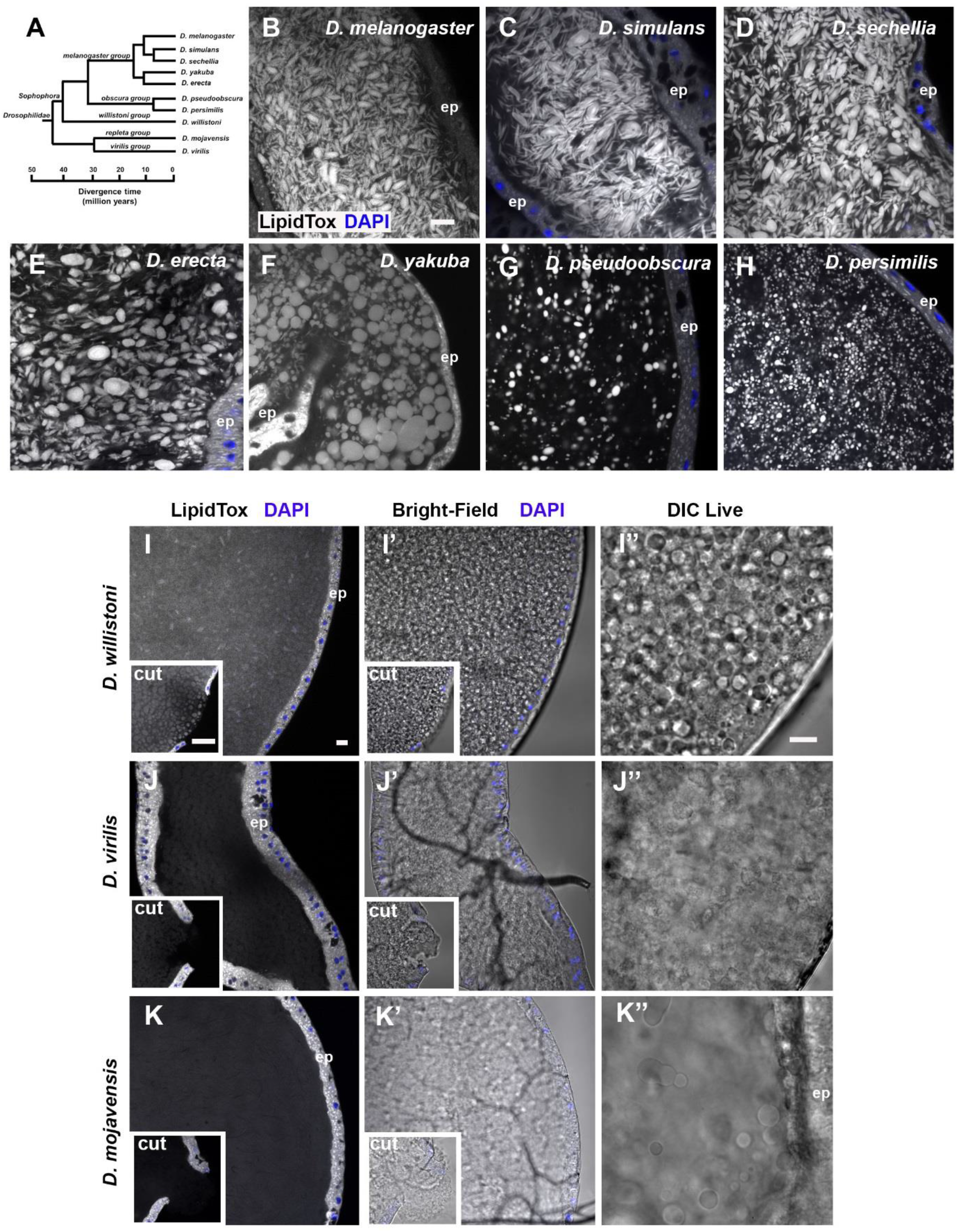
Co-evolution of microcarrier morphology and SP in *Drosophila* species. (A) Phylogenetic tree of *Drosophila* species used in this study. All species except *D. mojavensis* have a putative SP homologue. Adapted from http://flybase.org/blast/. (B-K) LipidTox staining of AGs from 6-day-old virgin males from selected *Drosophila* species, namely *D. melanogaster* (B), *D. simulans* (C), *D. sechellia* (D), *D. erecta* (E), *D. yakuba* (F), *D. pseudoobscura* (G), *D. persimilis* (*H*), *D. willistoni* (*I*), *D. virilis* (*J*) and *D. mojavensis* (K). For (I-K), where LipidTox microcarriers are not readily detectable, bright-field images of the same glands are shown (I’-K’), as well as DIC images (I’’-K’’) of different glands. Insets in (I, J, K) are images of AG with epithelial layer punctured to fully expose luminal contents to LipidTox stain, revealing stained structures only in *D. willistoni*. Note that different subgroups have noticeably different microcarrier size, shape and density. Nuclei marked with DAPI (blue). AG epithelium (ep). All scale bars, 10 μm; scale bar in (B) applies to (B-H); in (I), applies to (I-K) and (I’-K’); and in (I’’) applies to (I’’-K’’).

Examining more distantly related *Drosophila* species with more diverged SP proteins (Fig. S4) revealed very different microcarrier organisation. *D. pseudoobscura* and *D. persimilis*, both members of the *obscura* group, have smaller spherical microcarriers that appear to be more widely separated (Fig. 6G, H).

Two further *Drosophila* species *D. willistoni* and *D. virilis* express forms of SP with major structural differences to the *melanogaster* and *obscura* groups. These proteins not only lack a central 10 amino acid portion of the molecule, including the sequence required for proteolytic cleavage, but have also diverged considerably in other regions with the exception of the last 15 C-terminal amino acids (Fig. S4). Bright-field and DIC microscopy revealed *D. willistoni* glands have densely packed globular structures that can be stained with LipidTox, although this is most clearly observed in punctured glands (Fig. 6I-I’’). The *D. virilis* AG, whether intact or cut, showed little evidence of LipidTox-stained microcarriers (Fig. 6J) and bright-field and DIC imaging suggested a more uniform “flocculence” (Fig. 6J’, J’’). Finally, we examined the AGs of *D. mojavensis*, a species that lacks an SP homologue (Fig. S4). Although there were some large spherical structures in the gland lumen in DIC images, no LipidTox-positive staining was observed in these glands (Fig. 6K-K’). Therefore, our analysis suggests that the divergence of SP structure closely parallels changes in microcarrier shape, size and density.

## Discussion

Seminal fluid plays an essential role in male reproductive success. In *D. melanogaster*, SP, produced from the male AG, has been highlighted as a central player in this process, acting via receptors in the female to stimulate changes that increase fecundity and prevent remating. Here we demonstrate that SP has an additional, unsuspected role in males in the assembly of neutral lipid-containing microcarriers in the AG lumen (summarised in Fig. 5I). These microcarriers store SP, other proteins with lipid anchors or potentially hydrophobic domains, and neutral lipids, so that they can be delivered to females during mating, and dispersed rapidly in the female reproductive tract. Our analysis of microcarriers in other *Drosophila* species reveals that SP’s microcarrier assembly function may exist in species in which SP has more limited roles in modulating the PMR, suggesting that this function might have been critical in the evolution of this molecule.

### Lipid microcarriers provide a store, delivery vehicle and dispersal machinery for a subset of seminal proteins

Seminal proteins are produced throughout adult life, but these proteins are only transferred to females sporadically. Some of these proteins are then rapidly activated via mechanisms that are thought to include proteolytic cleavage and pH changes in the female reproductive tract (discussed in (5)). Our data suggest that microcarriers could contribute to this activation process. They are repositories for main cell-derived seminal proteins, which presumably partition from the aqueous phase of the AG’s secretions, either because of their lipophilicity or because they have binding partners on the microcarrier surface. In the male, molecules like SP bind specifically to microcarriers and not to AG epithelial cells, strongly suggesting that these surfaces are structurally distinct. Subsequent microcarrier dissipation in the female reproductive tract provides a mechanism for dispersing proteins like SP, so they can associate with receptors and cell membranes following mating.

Although both staining of normal microcarriers with lipophilic dyes and the homogeneous internal structure of large defective *SP^0^/Df(SP)* null ‘microcarriers’ observed with DIC strongly suggest that neutral lipids are a major component of these structures, their precise composition remains unclear. In addition, their non-spherical shape in wild type males suggests that architectural proteins are highly likely to be involved in establishing their final structure, a proposal supported by the *SP* mutant phenotype. It will now be important to identify these other structural constituents and to establish whether any of these, unlike SP, play evolutionarily conserved roles in seminal fluid production outside the *Drosophilidae* family.

Analysis of transcriptomics data from adult *Drosophila* organs reveals high level expression in the AG of multiple lipases that are predicted to be secreted (eg., CG5162, CG11598, CG11600, CG11608, CG13034, CG18258, CG18284, CG31872, CG34447; (37), (38), with all having been detected in proteomics analyses of seminal fluid (39, 8). These include proteins sharing homology with triacylglycerol lipases (eg., CG5162, CG13034, CG18258, CG34447). These lipases provide a potential mechanism to break down neutral lipid transferred in microcarriers to females, so the products can be used as fuel. Mammalian seminal fluid also contains lipases (40, 41, 42) and triacylglycerides (43, 44), suggesting that the latter may be required, perhaps as a male-derived nutrient source, in the reproductive system of all higher organisms.

Our identification of extracellular neutral lipid microcarriers as accessible stores of specific seminal proteins is reminiscent of the role of intracellular lipid droplets in storage of cytoplasmic and nuclear proteins (45, 46). Lipid droplets are able to dock with specific intracellular organelles to mediate their functions and deliver their cargos. It will be interesting to investigate whether the remnants of microcarriers, such as the microdomains observed with SP-GFP, are in any way targeted to specific cells or structures after transfer to females, as these storage vehicles break down.

It has previously been reported in *Drosophila* that males can adaptively modulate the relative balance of seminal proteins, including SP, in the ejaculate, depending on female mating status and the presence of rival males (47, 48). Loading of selected proteins on to microcarriers might provide a simple mechanism to control such rapid changes, if the transfer of these large structures can be differentially regulated compared to soluble proteins, for example by controlling the opening of the sphincters through which seminal fluid passes from the AGs to the ejaculatory duct.

### Regulation of microcarrier morphology by SP and microcarrier/SP co-evolution in *Drosophila*

Our study reveals a previously unsuspected male-specific, SPR-independent role for SP in regulating microcarrier shape and size. *SP* mutants in *D. melanogaster* still have neutral lipid-containing structures, but they appear to aggregate and fuse, particularly after mating, to generate large lipid droplet-like structures that no longer retain molecules like SP at their surface. To date, we have not been able to separate the different activities of SP in males and females through expression of different mutants or altered SP levels, making it difficult to fully gauge the importance of the male-specific microcarrier function. However, the observation that *SP* mutants, which were known to affect binding of SP to the surface of sperm or its subsequent release, also fail to rescue the microcarrier defect in *SP* null males, suggests that the interpretation of the phenotypes associated with these mutants requires some re-evaluation.

Tsuda et al. (24) have suggested that SP is likely to have roles in addition to its effects mediated via SPR signalling in the female reproductive tract, which include induction of a female sexual refractory period. This is because some SP-expressing species like *D. pseudoobscura* and *D. persimilis* do not appear to express SPR in this location and additionally show much female less post-mating refractoriness relative to other SP-producing species (49). Our data (Fig. 6) suggest that microcarrier assembly may be this additional function with the shape of microcarriers rapidly co-evolving with SP, and an absence of microcarriers in species with a highly divergent (*D. virilis*) or no (*D. mojavensis*) SP homologue. It will be interesting to investigate whether other proteins with fundamental roles in packaging and storing seminal fluid components have also evolved signalling roles in some species.

An important conclusion from our study is that the normal transfer of several different seminal proteins is likely to be interdependent. Elegant studies by the Wolfner lab have identified several long-term response (LTR) network genes expressed in the AG that are required either in the male or female for SP to be retained in the sperm storage organs (50, 51, 52). It will be important to investigate whether any of these genes is involved in loading or unloading SP from microcarriers, or indeed, whether they play a role in microcarrier assembly. Furthermore, determining whether other Acps or main cell-expressed GPI-anchored proteins are microcarrier cargos should allow the functions of these structures to be assessed more extensively and may suggest molecular tools that could be used to screen for similar processes in higher organisms.

## Materials and Methods

### *Drosophila* Stocks and Genetics

Fly stocks were obtained from the following sources: the Bloomington *Drosophila* Stock Center provided *UAS-GFP.nls* (53)3, *UAS-mCD8-GFP* (54), *tub-GAL80^ts^* (55), *UAS-SP-RNAi#2* TRiP.JF02022 (56), *UAS-mCD8-ChRFP*; the Vienna *Drosophila* Resource Center provided *UAS-SP-RNAi#3* (v109175); the Kyoto DGRC Stock Center provided *spi-GAL4* (57); S. Goodwin provided *dsx-GAL4* (58), *Acp26Aa-GAL4* (11), *SP-GFP* (33); T. Aigaki provided *UAS-SPn-GFP-SPc, UAS-SPn-GFP, UAS-GFP-SPc, UAS-sGFP* (24); M. Wolfner provided *SP^QQ^*, *SP^Δ2-7^* (22), *Df(SPR)* (25); S. Eaton provided *UAS-GFP-GPI* (59); T. Chapman provided *SP*^0^, *SP*^0^ *SP*^+^ (12), *Df(3L)Δ130* (34), *UAS-SP-RNAi-IR2* (RNAi#1; (11)); L. Partridge provided *w^1118^*. A. McGregor provided *D. simulans, D. sechellia, D. yakuba, D. pseudoobscura, D. virilis* and the Gulbenkian Institute provided *D. erecta, D. persimilis, D. mojavensis.*

### Fly husbandry

Flies were maintained on standard cornmeal agar food (12.5 g agar, 75 g cornmeal, 93 g glucose, 31.5 g inactivated yeast, 8.6 g potassium sodium tartrate, 0.7 g calcium, and 2.5 g Nipagen [dissolved in 12 ml ethanol] per litre) at 25°C on a 12:12-h light:dark cycle. Males for *SP* knockdown or those with *tub-GAL80^ts^* were shifted to 29°C on eclosion to activate UAS-transgenes.

### Staining and immunostaining of fly reproductive tracts

Unless otherwise stated, 3-4-day-old virgin males were used for AG dissection and for mating experiments. 4-7-day-old *w^1118^* virgin females were used for mating experiments. For fixed tissues, reproductive tracts were dissected in 4% paraformaldehyde (Sigma-Aldrich) in PBS (Gibco). AGs with the ejaculatory duct attached were fixed for 20 min and rinsed at least four times in PBS prior to further treatments. For females the abdomen was carefully opened up and fixative allowed to permeate internally for 20 min prior to removal of the uterus with seminal receptacle, spermathecae and common oviduct attached. Reproductive tracts were washed four times with PBS.

Fixed accessory glands were stained at room temperature in the following solutions and then washed four times in PBS: 1:50 dilution in PBS of a 10 mg/ml solution of Nile red (Sigma-Aldrich) dissolved in acetone and incubated for 30 min; 1:100 dilution in PBS of LysoTracker Deep Red (Life Technologies) for 1 h; 1:50 dilution in PBS of LipidTox (Invitrogen) for 1 h; 1:40 dilution in diluent C of a 1mM stock of PKH26 red fluorescent cell marker (Sigma-Aldrich) for 30 min; 1:1000 dilution in PBS of a 10 mg/ml stock of Hoechst 33342 (Invitrogen) for 5 min.

For live imaging, accessory glands were dissected in ice-cold PBS. Live glands requiring staining were treated for 5 min in a 1:100 dilution of LysoTracker Red DND-99 (Life Technologies) in ice-cold PBS.

For ANCE antibody staining, fixed accessory glands were permeabilised for 6 x 10 min in PBST (1 X PBS, 0.3% Triton X-100 [Sigma-Aldrich]), blocked for 30 min in PBSTG (PBST, 10% goat serum [Sigma-Aldrich]) and incubated overnight at 4°C in rabbit anti-ANCE primary antibody (60) diluted 1:2000 in PBSTG. Glands were then washed for 6 x 10 min in PBST before incubation in a 1:400 dilution of Cy-5-conjugated donkey anti-rabbit secondary antibody (Jackson Laboratories) for 2 h at room temperature. Glands were further washed in PBST for 6 x 10 min prior to mounting.

Glands stained with Hoechst were mounted in PBS; all other fixed reproductive tracts were mounted in Vectashield with DAPI (Vector Laboratories). Glands for live imaging were mounted in a small drop of ice-cold PBS surrounded by 10S Voltalef (VWR) halocarbon oil (61).

### Electron microscopy

3-day-old *w^1118^* male reproductive tracts were dissected and incubated overnight in 2.5% glutaraldehyde and 4% formaldehyde in PBS (pH 7.2). Glands were then washed with PBS, refixed in 1% osmium tetroxide (Agar Scientific) for 20 minutes, washed 3 times in distilled water and dehydrated through a graded alcohol series and incubated in ethanol and Spurr’s epoxy resin (1:1) (Agar Scientific). Glands were embedded in 100% Spurr’s epoxy resin between two sheets of polythene and polymerised overnight at 60°C. Ultrathin sections were prepared with a Reichert Ultracut R Ultramicrotome (Leica Biosystems) and mounted on formvar-coated slot grids (Agar Scientific). Sections were stained with 2% uranyl acetate and lead citrate (Agar Scientific), and imaged using a JEOL 1010 electron microscope (80kV).

### Imaging

Images of fixed reproductive tracts were acquired either on a Zeiss LSM 510 Meta [Axioplan2] or a LSM 880 laser scanning confocal microscope equipped with Zeiss 10x NA 0.45, 20x NA 0.8, 40x NA 1.3 and 63x NA 1.4 objectives. Live scanning confocal imaging was performed on a Zeiss LSM 710 microscope using a 63x NA 1.4 objective. Live DIC images were acquired on a DeltaVision Elite wide-field fluorescence deconvolution microscope (GE Healthcare Life Sciences) equipped with a 100x, NA 1.4 UPlanSApo oil objective (Olympus).

### Automated analysis of microcarrier size

Images were opened with Fiji software. Microcarrier image analysis was performed using the open‐access CellProfiler Software version 2.2.0. A workflow for segmenting all the microcarriers and measuring the minimum feret diameter of each microcarrier was developed by adding pre-programmed algorithmic modules in a pipeline. Histograms based on microcarrier minimum width and microcarrier area in different minimum width ranges were plotted using GraphPad Prism-8 software.

Changes in microcarrier size were further assessed by recording the presence or absence of microcarriers with a minimum width greater than 10 μm for 7-10 glands in a representative 100 μm^2^ field of view of the lumen midway along the length of the gland. *P*-values were calculated using Fisher’s exact test.

All materials, tools and datasets generated in this study are presented in the paper or will be made available upon request.

## ACKNOWLEDGEMENTS

We thank Suzanne Eaton, Mariana Wolfner, Toshiro Aigaki, Tracey Chapman, Stephen Goodwin, Elwyn Isaac, Nuno Soares and Alistair McGregor for stocks and reagents; we are grateful to the Bloomington *Drosophila* Stock Center, the Vienna *Drosophila* Resource Center and the Kyoto DGRC Stock Centre for flies. We thank the Micron Advanced Bioimaging Unit (supported by Wellcome Strategic Awards 091911/B/10/Z and 107457/Z/15/Z) for their support & assistance in this work. We acknowledge the support of the Biotechnology and Biological Sciences Research Council (BB/K017462/1, BB/L007096/1, BB/N016300/1, BB/R004862/1 and a Fellowship to SW, BB/K014544/1), Cancer Research UK (C19591/A19076, C602/A18974), the Cancer Research UK Oxford Centre Development Fund (C38302/A12278), and the Wellcome Trust (Strategic Awards #091911, #107457 and 102347/Z/13/Z), as well as the MRC for studentship funding.

## Supplementary Figures

**Fig. S1.**
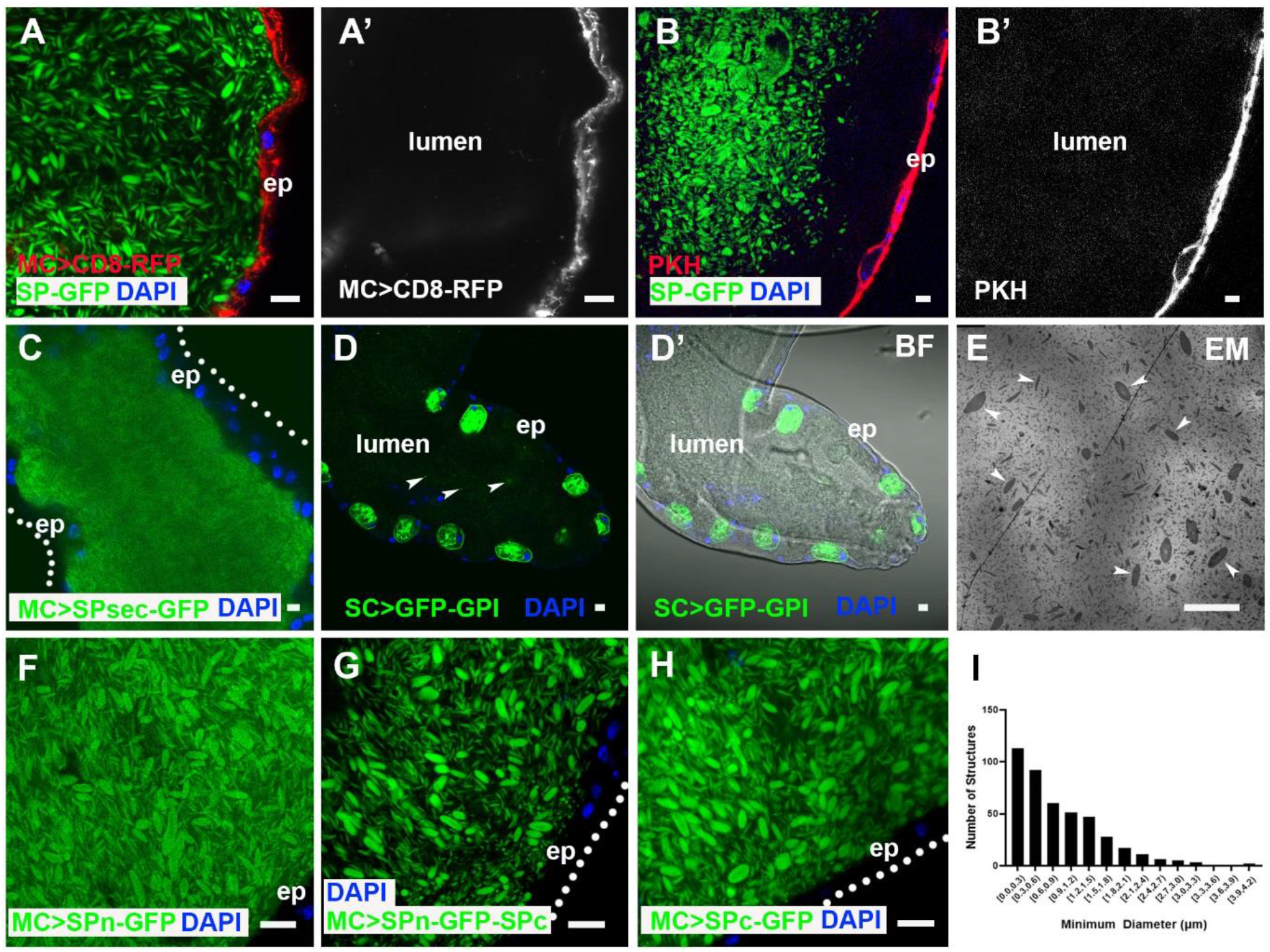
Structure and cargos of accessory gland microcarriers. (A, B) Both main cell-expressed transmembrane CD8-RFP (A) and the lipid bilayer dye PKH26 (B) mark the main cell apical membrane (A’, B’), but not the luminal microcarriers labelled by SP-GFP (merge in A, B). (C) Secreted GFP is not preferentially partitioned on to microcarriers. (D) In the AG lumen, secreted secondary cell-expressed GFP-GPI is primarily associated with puncta (arrowheads; cell boundaries seen in bright-field image; D’) and does not appear to be loaded on to microcarriers. (E) Transmission electron micrograph of AG lumen showing a range of sizes of microcarriers (arrowheads) (F-H). The N-terminus of mature SP with a C-terminal GFP tag (F; SPn-GFP), full length SP with a central GFP tag (G; SPn-GFP-SPc) and the C-terminus of SP with an N-terminal GFP tag (H; GFP-SPc) all concentrate on microcarriers when expressed in main cells albeit at lower levels for SPn-GFP. (I) Size distribution of microcarriers in LipidTox-stained AG lumen from image in Fig. 1B’. Nuclei marked with DAPI (blue). AG epithelium (ep). Scale bars, 10 μm.

**Fig. S2.**
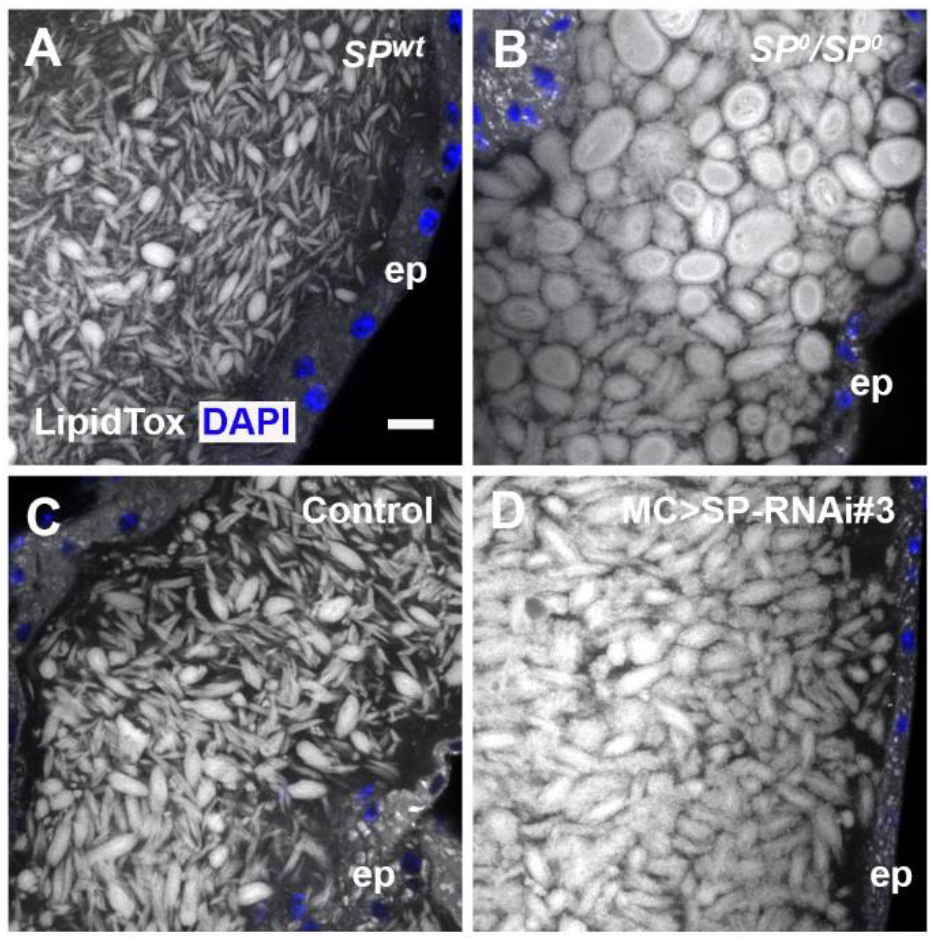
Loss or reduction of SP disrupts microcarrier morphology. (A-D) LipidTox-stained microcarriers are highly enlarged in *SP^0^/SP^0^* homozygous males (B) when compared to controls (A). Expressing a third RNAi targeting *SP* transcripts *(UAS-SP-RNAi#3*; v109175) induces the formation of enlarged microcarriers (D), unlike controls (C). Nuclei marked with DAPI (blue). AG epithelium (ep). Scale bars, 10 μm.

**Fig. S3.**
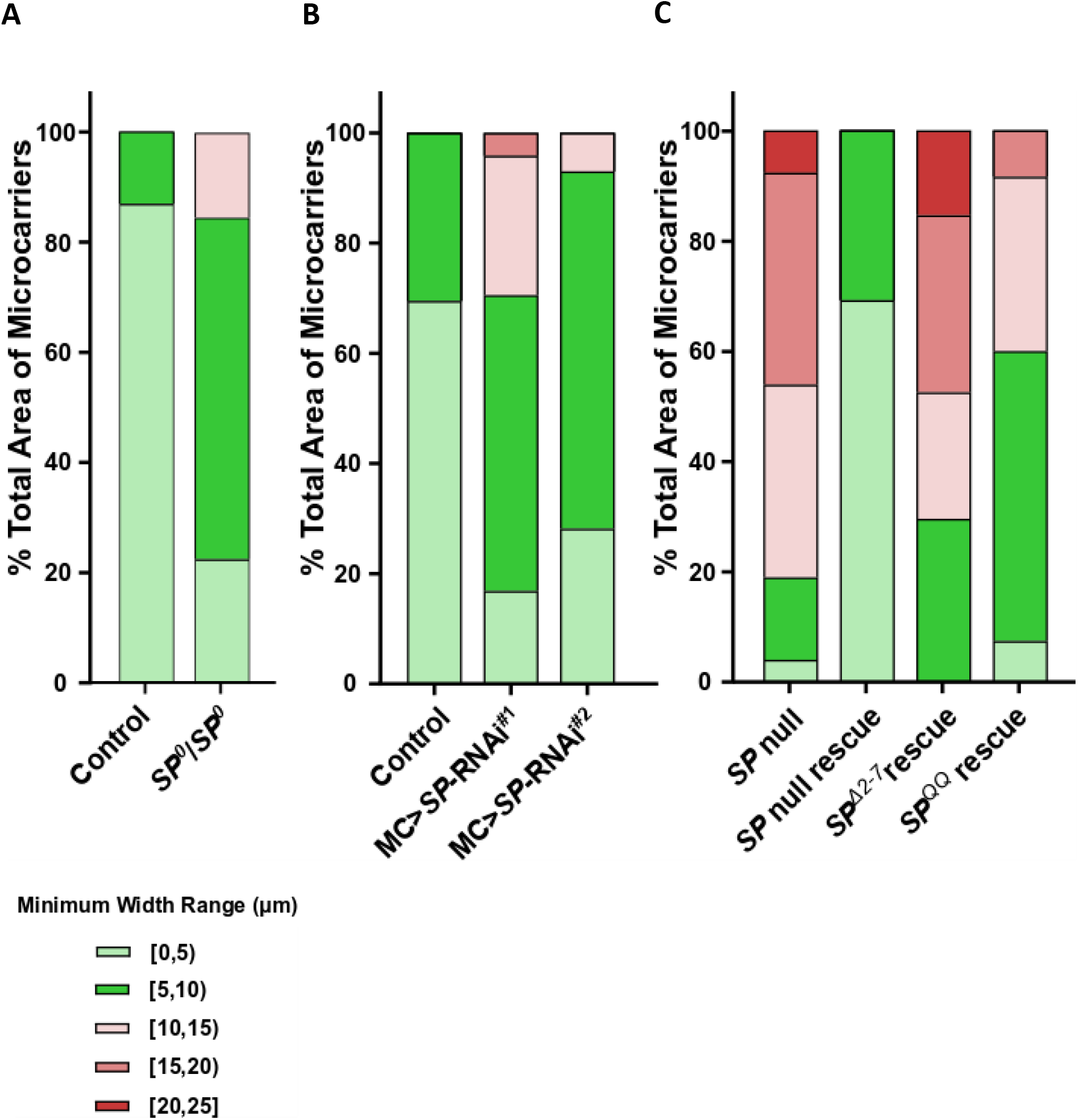
*SP* mutant, *SP* RNAi and some *SP* rescue glands have enlarged microcarriers. (A-C) Histograms showing percentage of microcarrier area within given microcarrier minimum width ranges. (A) More lipid is incorporated into microcarriers in larger size ranges in homozygous *SP^0^/SP^0^* gland (from Fig. S2B) than in control glands (Fig. S2A). (B) *SP*-RNAi knockdown in main cells increases the incorporation of lipid into larger microcarriers (from glands shown in Fig. 4B, C) compared with control glands (Fig. 4A). (C) Expression of SP^Δ2-7^ or SP^QQ^ (from glands shown in Fig. 4J, K) fails to rescue the *SP^0^/Df(SP)* null phenotype (Fig 4H) when compared with *SP^0^ SP^+^/Df(SP)* rescue (Fig. 4I). Main cell (MC).

**Fig. S4.**
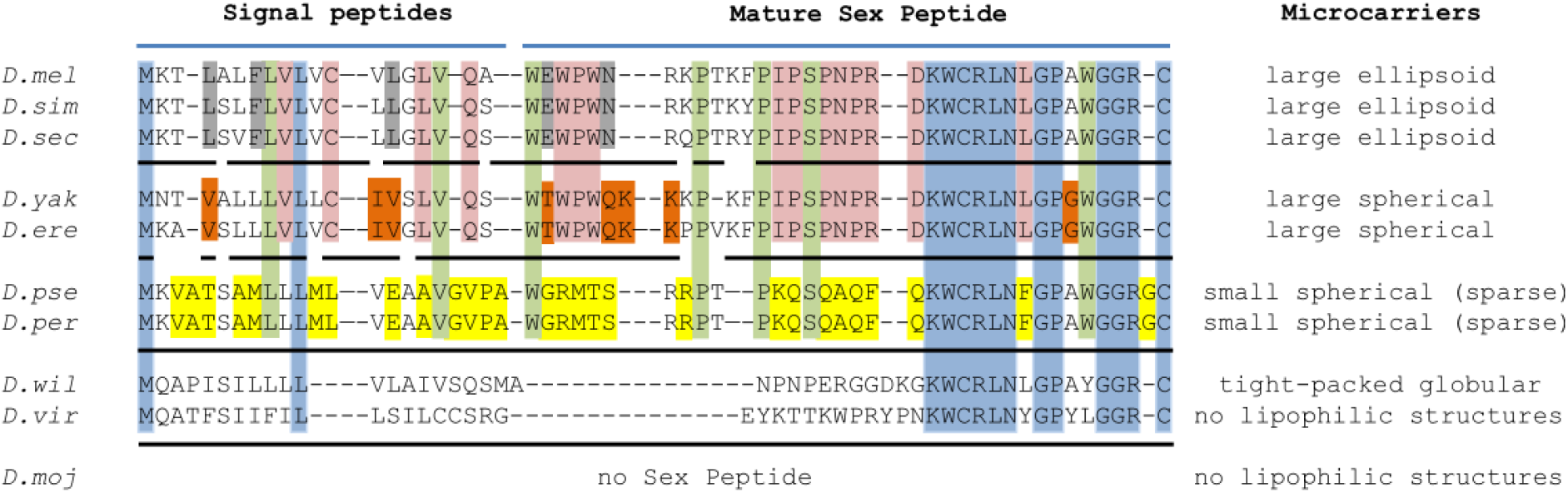
Rapid evolution of SP sequences correlates with changes in microcarrier morphology, size and abundance in diverse *Drosophila* species. SP protein sequence alignment for the different *Drosophila* species used in this study. Species are clustered according to subgroup. Conserved amino acids are highlighted or underlined; blue = conserved across all SP-expressing species; green = conserved in *melanogaster* and *obscura* groups; pink = conserved in *melanogaster* group; grey, brown or yellow = conserved within a single species cluster. The subdivision of these groups correlates with SP protein sequence, and shape, size and abundance of microcarriers in each species. Adapted from (24).

**SI Movie 1. Live imaging of SP-GFP-labelled microcarriers.**

*Z* stack through distal tip of live SP-GFP AG. Lysotracker-Red (red; used at high concentrations) marks epithelial layer of gland. Confocal sections were captured at 1.5 μm intervals on *Z* axis.

**Supporting information Movie 1.**

**Live imaging: *Z* stack through distal tip of SP-GFP AG.**

